# The evolution of red blood cell shape in a continental radiation of fishes

**DOI:** 10.1101/2020.04.03.023994

**Authors:** Brenda Oliveira Martins, Lilian Franco-Belussi, Mayara Schueroff Siqueira, Carlos E. Fernandes, Diogo B. Provete

**Author notes:** These authors contributed equally to this work (Brenda Oliveira Martins, Lilian Franco-Belussi). Author contributions: BOM collected cell morphometric data assisted by MSS, LFB. CEF and DBP conceived the study. DBP analysed and curated the data. DBP, LFB, and BOM prepared the first draft of the manuscript to which all authors contributed afterwards. Corresponding author: Diogo B. Provete, Instituto de Biociências - INBIO, Universidade Federal de Mato Grosso do Sul – UFMS, Av. Costa e Silva, s/n, Bairro, Universitário, Campo Grande, 79002-970, Mato Grosso do Sul, Brasil.

## Abstract

The size and shape of Red Blood Cells (RBC) can provide key information on life history strategies in vertebrates. However, little is known about how RBC shape evolved in response to environmental factors and the role of phylogenetic relationship. Here, we analyzed RBC morphometrics in a continental radiation of fishes testing the hypothesis that phylogenetic relationship determines species occupation of morphospace. We collected blood samples of five specimens of 15 freshwater fish species from six orders and used basic stereological methods to measure cell and nucleus area, perimeter, and diameter, cell and nucleus volume, nucleus:cytoplasm ratio, and shape factor of 50 cells per specimen. Then, we conducted a phylogenetic Principal Components Analysis using a dated phylogeny and built a phylomorphospace. To test if the phylogenetic relationship predicted the phenotypic similarity of species, we calculated multivariate phylogenetic signal. We also estimated the evolution rate of RBC shape for each node and tip using ridge regression. Finally, we tested if the position in the water column influenced RBC shape using a phylogenetic GLS. RBC shape seems to have evolved in a non-stationary way because the distribution pattern of species in the phylomorphospace is independent of the phylogeny. Accordingly, the rate of evolution for shape was highly heterogeneous, with an increase in the genus *Pygocentrus*. Water column position does not influence RBC shape. In conclusion, RBC shape seem to have evolved in response to multiple selective pressures independent of life history characters.

## Introduction

The shape, size, and number of Red Blood Cells (RBC) vary greatly in vertebrates (Wintrobe 1934). Although the reasons for this variation are not well understood, phylogenetic relationships can at least partially explain it, because early diverging vertebrate lineages have larger but fewer RBC (Anderson et al., 2018). Early diverging vertebrates also have round and flattened RBCs with a central nucleus, while derived lineages have ellipsoid cells (Nikinmaa 1990; Glomski et al. 1992).

Both abiotic factors and life history traits can also affect the composition and morphology of fish RBC (Dal’Bó et al., 2015; Kumar, 2016; Yakhnenko et al., 2016). For example, RBC size is inversely related to swimming ability in teleosts (Lay and Baldwin 1999; Witeska 2013), while size of RBCs is highest at intermediate altitude in one lizard species (González-Morales et al. 2017). In addition, cell size has an inverse relation with metabolic rate, suggesting lowest metabolic cost to maintain active membrane transport in large cells (Szarki, 1970; Graham et al. 1985, Maciak et al, 2011). Deep-sea fish species have large RBC, compatible with their low metabolism (Graham et al. 1985). Then, RBC size in teleost has an adaptative value and changes in response to external factors (Lay and Baldwin 1999). RBC volume is another morphological variable correlated with physiological state in fish. For example, cell volume is negative correlated with hemoglobin concentration, while nuclear volume explains only 17% of cell volume (Lay and Baldwin 1999).

The nucleus of RBC is strongly related to species adaptation to environmental conditions. Nucleus size is positively correlated with genome size in vertebrates (Gregory 2001). Accordingly, the amount of DNA regulates the size of cells as well as the nucleus. Larger nuclei have more room for protein synthesis that are required in the metabolism of larger cells (Gregory 2001). Variations in RBC DNA content are also directly related to hematopoiesis, which in turn influence species adaptation. Genome size seems also related to life-history traits. For example, genome size is positively related with the size and diameter of eggs, and negatively related with growth rate and standard size in fish (Hardie and Hebert, 2004; Smith and Gregory 2009).

The shape of RBC also changes according to cell maturation stage. While immature RBC (erythroblasts) are usually rounded, older cells are fusiform (Fijan 2002, Passantino et al. 2004). However, the degree of ellipticalness seems to vary among species (Dal’Bó et al., 2015). However, little is known about the physiological implications of the variation in the fusiform shape for species. Similarly, little is known about the factors involved in the variation of RBC volume and its implications (Lay and Bladwin 1999) but increasing temperature and body mass increase RBC volume in tetrapods (Gillooly and Zenil-Ferguson 2014). Also, RBC volume is related to polyploidy in fish (Gregory 2001). However, little is known about the relative role of phylogenetic relationships and abiotic factors in determining RBC shape.

No study has yet analysed how RBC shape have evolved in fish using modern phylogenetic comparative methods. As shape is multivariate, here we used a set of first- and second-order stereological variables to describe the shape of RBC. Also, we tested if the position in the water column, as a proxy for partial oxygen pressure, species are usually found affect RBC shape. Our hypothesis is that neustonic species (that live closer to the surface) would have smaller RBC due to higher surface area to volume ratio. We also built the phylomorphospace of RBC shape and tested for phylogenetic signal. We expect species with similar habitat use would be close together in the morphospace. This is the first study combining common stereological analysis and phylogenetic comparative method to understand how fish RBC evolved. Our results can help understand how fish species evolved to occupy habitats with different oxygen concentration or toxicity, besides understanding how metabolic rate vary among species.

## Methods

### Specimen sampling

We analyzed five individuals of 15 species belonging to 15 genera and 2 superorders: *Pygocentrus nattereri, Gymnotus inaequilabiatus, Astyanax lineatus, Bujurquina vittata, Cichlasoma dimerus, Metynnis maculatus, Prochilodus lineatus, Piaractus mesopotamicus, Rhamdia quelen, Poecilia reticulata, Hypostomus boulengeri, Pseudoplatystoma corruscans, Synbranchus marmoratus, Brycon hilarii*, and *Hyphessobrycon anisitsi* (Figure 1). We obtained adult fish specimens from both sexes between 2017 and 2018 from a number of sources: *Pygocentrus nattereri* was fished in a river the Pantanal field station (Corumbá, Mato Grosso do Sul, central Brazil); *Gymnotus inaequilabiatus, Hyphessobrycon anisitsi*, and *Synbranchus marmoratus* were bought from a fish store; *Astyanax lineatus* and *Bujurquina vittata* were collected in an urban lake in Campo Grande; *Cichlasoma dimerus, Metynnis maculatus, Prochilodus lineatus, Piaractus mesopotamicus* were donated by the city aquarium; *Pseudoplatystoma corruscans*, Brycon hilarii, and *Hypostomus boulengeri* were obtained from the university Aquaculture Research ponds; *Poecilia reticulata* was collected in an urban stream in Campo Grande. These species were selected because they are common in our region, use distinct habitats, and belong to distant lineages.

**Figure 1.**
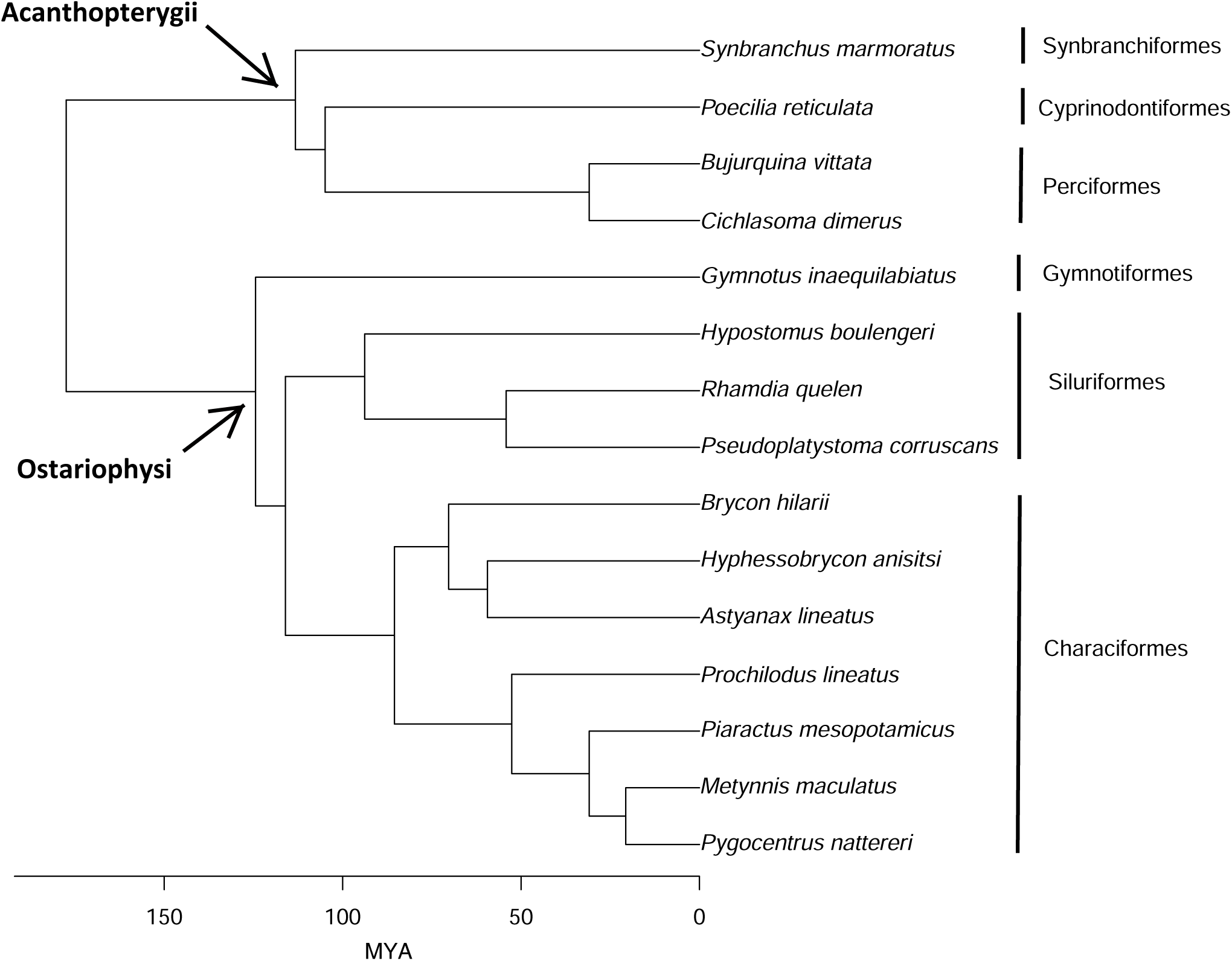
Dated phylogenetic tree with species to which we have phenotypic data used in the analysis, pruned from Rabosky et al. (2018).

Fish were anaesthetized with eugenol solution (50 mg L^-1^) before blood sampling. Blood was collected by caudal vein or cardiac puncture with syringe with 3% EDTA or by decapitation in small species (Ranzani-Paiva et al., 2013). Blood smears were stained with May-Grunwald-Giemsa-Wright for erythrocyte morphometry (Tavares-Dias and Moraes 2002). Detailed protocol is available at Rodrigues et al. (2020; see also Figure S1).

### Red blood cell stereology

Design-based stereological methods have been used since the 1970s to quantify shape and size changes in cells and histological structures (West 2012). This approach uses linear (first-order) measurements to derive tridimensional (second-order) variables that describe the shape of cells in tissues. To take measurements, we used a digital camera (Nikon D3500) coupled in a light microscopy (Zeiss Primo Star^®^). Digital images in 1000x magnification were used to measure the following first-order variables: area (µm^2^), perimeter, and largest and smallest diameter (µm) of cells and area, perimeter, and diameter of the nucleus in 50 randomly selected cells per animal (see Supporting material), i.e., 250 cells per species (50 cells * 5 animals). From these linear measurements, we calculated the following second-order stereological measurements: To calculate length, we took the mean of the largest and smallest diameters. Then, to calculate radius, we simply divided length by 2. Cell volume (µm^3^) was calculated using the formula: 

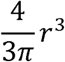

where *r* is radius (see West 2012); nucleus volume (µm^3^) was calculated by the formula: 

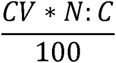

where CV is cell volume and N:C is nucleus:cytoplasm ratio. Cell and nucleus circularity (or shape factor) were estimated by the formula: 

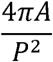

where *A* is cell area and *P* cell perimeter (Russ and Dehoff 2000). Circularity ranges from 0 (elliptical shape) to 1 (circular shape). All measurements were made in Motic Images Plus 2.0 in a double-blind fashion (see Fig. S2).

We calculated the mean of each variable for each species (Table S1), which were used as phenotypic traits in further analysis. To visualize the correlation pattern between variables, we built a correlogram considering the average of each variable (Fig. S3) using the *corrplot* package (Wei and Simko, 2017) of R software.

### Phylogenetic comparative methods

To carry out the comparative analyzes, we first obtained a phylogeny with branch length in millions of years representing the relationship of the 15 species for which we have phenotypic data (Figure 1). This phylogeny was obtained from a fully-sampled topology for Actinopterygii recently published (Rabosky et al., 2018). Analysis was done in the fishtree package (Chang et al., 2019).

Then, we performed a phylogenetic Principal Components Analysis (pPCA) on the matrix of phenotypic data with the eight variables considering the phylogeny (Revell, 2009). The pPCA was performed using the correlation matrix of variables and assuming that traits have evolved according to a Brownian Motion (BM) model of evolution. The pPCA uses a **C** matrix that describes the variance-covariance structure between species given by the phylogeny and the evolutionary model (BM in this case) to obtain the Principal Components and species scores (see also Adams & Collyer 2019). Analysis was performed using the `phyl.pca` function of the phytools package (Revell, 2012).

From this exploratory analysis, we built a morphospace for all 15 species showing their similarity in relation to RBC shape. The analysis estimates the node values using a Maximum Likelihood method. Finally, we built the phylomorphospace by projecting the phylogeny onto the morphospace built with the first two Principal Components, which allows us to explore how lineages occupied the morphospace as their traits evolved (Adams and Coller 2018, 2019). Analysis was performed using the `phylomorphospace` function of the phytools package.

Finally, we calculated a phylogenetic signal measure for multivariate data called K_mult_ (Adams, 2014) using all pPCs to avoid missing variation (Uyeda et al., 2015). This method allows calculating a statistic that measures how similar closely related species are in relation to their phenotypic measurements. Blomberg’s K (Blomberg et al., 2003) is a measure widely used to measure phylogenetic signal for univariate data and ranges from 0 <K> 1. High phylogenetic signal (K>1) indicates that closely related species tend to be similar in relation to phenotypic attributes (Münkemüller et al., 2012). This method has been adapted for multivariate data recently (Adams, 2014), along with a Monte Carlo randomization procedure that allows testing the significance of the statistic. Analysis was performed with the `physignal` function of the geomorph package (Adams et al. 2020).

To test the effect of the position in the water column species usually occupy on RBC shape, we gathered data on depth range from FishBase (Froese and Pauly 2019) and treated it as a categorical variable, henceforth referred to as habitat. Thus, species were classified as demersal (n=4), pelagic (n=2), and benthopelagic (n=9). Afterwards, we compared the fit of evolutionary models to the eight pPCs. Specifically, we fit Pagel’s lambda transformation, Brownian Motion, and Ornstein-Uhlenbeck models to data using the leave one out cross-validation of the penalized log-likelihood, as implemented in the `mvgls` function (Clavel and Morlon 2020) of the package mvMORPH (Clavel et al. 2015) including habitat as a predictor variable. Then, we compared the model fit using the Extended Information Criterion (EIC), which showed that the best model was Brownian Motion (BM; Table S2). We built a phylogenetic Generalized Least Squares (Clavel and Morlon 2020) using BM to test the effect of habitat on RBC shape, with the Pillai’s trace as test statistic and Type I Sum of Squares.

Finally, to test the hypothesis if a change in evolution rate is the mechanism allowing species to occupy different positions in the morphospace, we estimated tip-level evolution rate of RBC shape using phylogenetic ridge regression, as implemented in the RRphylo package (Castiglione et al. 2018). For this analysis we used all eight pPCs. This analysis also automatically finds nodes in which there was a rate shift. The R markdown dynamic document describing how analyses were conducted and associated data are available at FigShare (Martins et al. 2020).

## Results

The shape of Red Blood Cells (RBC) of fish species studied varied from oval to ellipsoid and a rounded nucleus (Figure 2). Cell size varied by an order of magnitude, from the largest cell of *Synbranchus* to the smallest in *Poecilia*. The same phenomenon happened to cell volume (Fig. 3). Nucleus area varied from 7.45 µm^2^ (± 1.2 SD) in *Poecilia* to 27.30 µm^2^ (*±* 4.9 SD) in *Synbranchus*. Additionally, the range of intraspecific variation in some measurements varied among species. For example, cell volume had the highest standard deviation, while shape factor, nucleus area and cell perimeter seem to vary less within species (Table S1).

**Figure 2.**
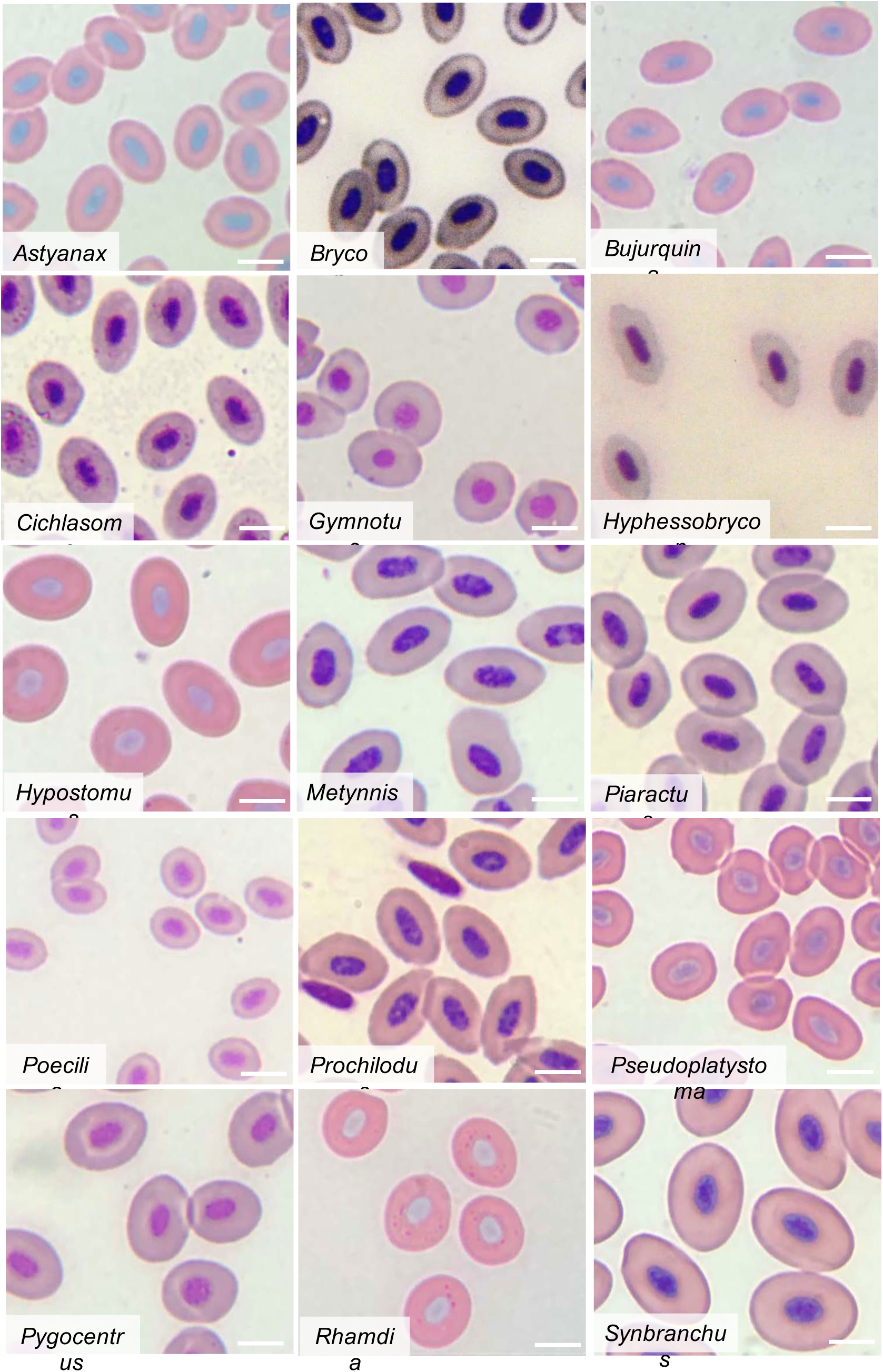
Shape and size of Red Blood Cells of the 15 fish species analysed. Staining: MMGW. Scale bars = 10 µm.

**Figure 3.**
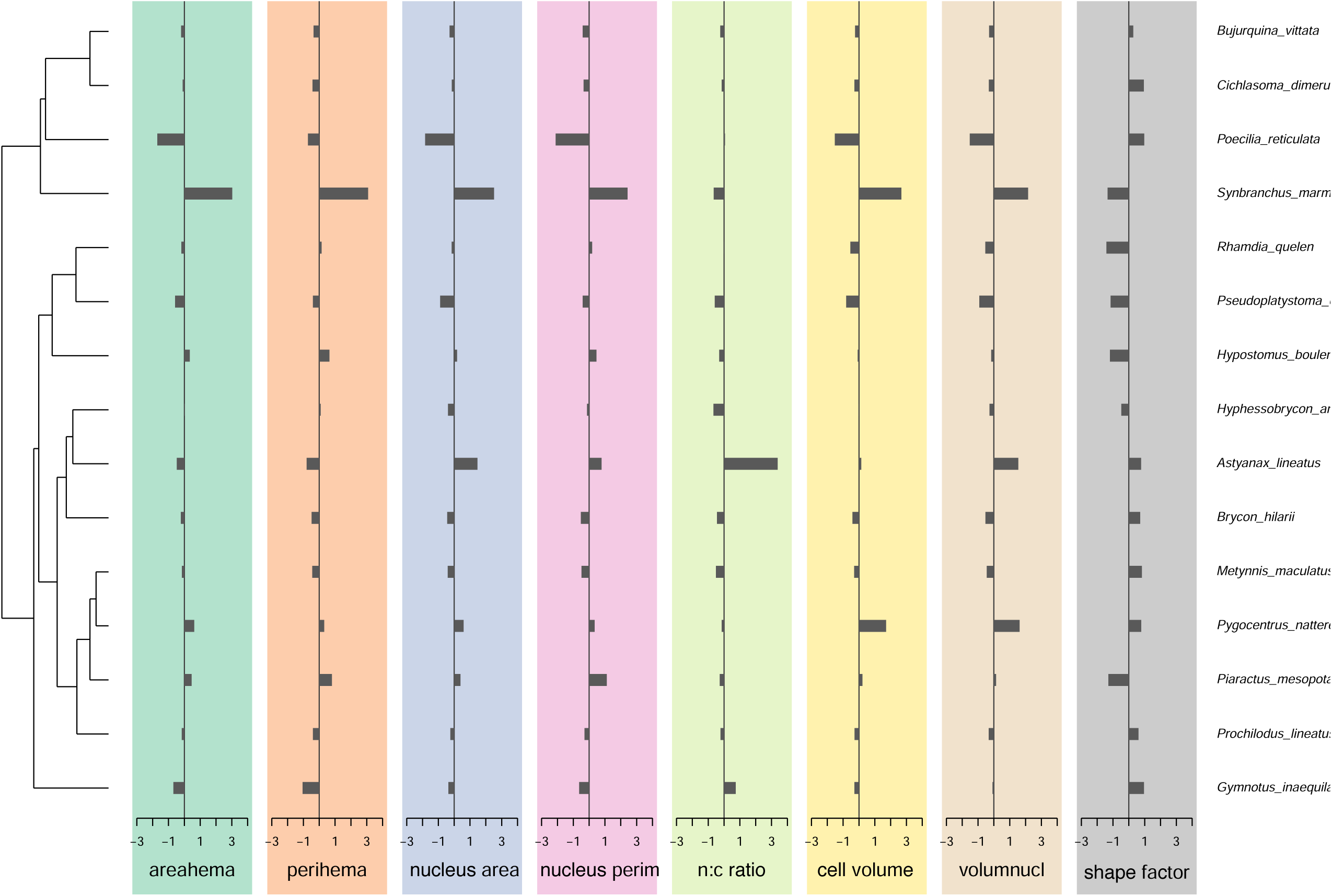
Diagram showing the mean of each stereological variable with the phylogenetic tree of the species sampled. Data on standard deviation for each variable is available at Table S1.

The pPC1 retained 66.1%, while pPC2 retained 21.4% of the variation in RBC shape. Cell and nucleus area, cell and nucleus perimeter, cell and nucleus volume were highly negatively correlated (> 0.84) with pPC1, whereas the nucleus:cytoplasm ratio was highly (−0.87) correlated with pPC2 (Table S2). Species positively correlated with pPC1 had higher shape factor, whereas those negatively correlated had higher cell and nucleus area, nuclear and cell perimeter, cell and nucleus volume, and nucleus:cytoplasm ratio (Table S2). Species positively correlated with pPC2 had higher cell perimeter, cell and nuclear area, whereas those negatively correlated had higher nucleus:cytoplasm ratio, nucleus and cell volume, and nucleus area (Fig. S4).

The distribution of species in the phylomorphospace suggest that closely related species do not have RBC with similar shape (Fig. 4). Accordingly, RBC shape does not exhibit phylogenetic signal (K_mult_ = 0.6317; Effect size = 0.7863; *P* = 0.159). The low effect size suggest phylogenetic signal is not concentrated in a few dimensions. To test if only a subset of dimensions displays phylogenetic signal (see Adams & Collyer 2019), we re-run the analysis with only the first two pPCs. Results did not change (see Martins et al. 2020). However, the morphospace suggest a non-stationary pattern in the evolution of RBC shape, since most closely related species occupy distant positions, but the position of siluriformes (*Pseudoplatistoma, Rhamdia*, and *Hypostomus*) mirror their phylogenetic relationship in the upper part of the morphospace. Interestingly, the Acanthopterygii included in our sampling: *Synbranchus, Poecilia, Bujurquina*, and *Cichlosoma* had the most divergent pattern, occupying extreme positions in the phylomorphospace, whereas species of Ostariophysi were more packed together. There is a high divergence in RBC shape within Characiformes. Therefore, there seems to be a variation in the pattern of RBC shape evolution at the level of higher order groups (Superorders), instead of a homogeneous pattern throughout the whole phylogeny.

**Figure 4.**
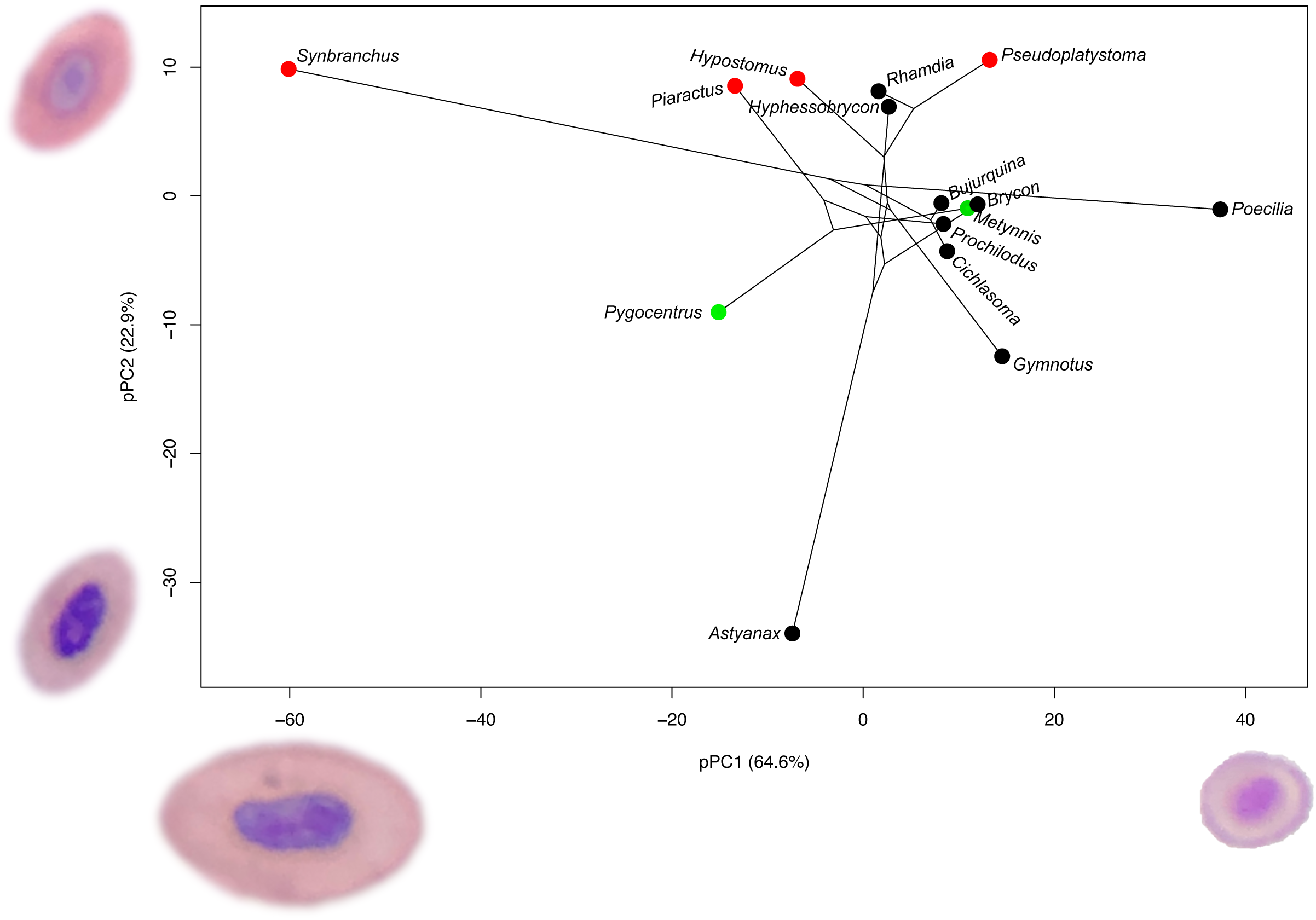
Phylomorphospace constructed with phylogenetic Principal Components 1 and 2 showing species distribution in the reduced space. Red Blood Cell (RBC) images are all in the same magnification and belong to species occupying the extreme positions along each pPC. The RBC associated positively with pPC1 is from *Poecilia*, which has a high shape factor, while the RBC associated negatively with pPC1 is from *Synbranchus* that has high cell area and perimeter, and high nucleus area and perimeter. The RBC associated positively with pPC2 belongs to *Pseudoplatystoma*, which has high cell perimeter. The RBC image associated negatively with pPC2 is from *Astyanax* and has a high shape factor, high nucleus:cytoplasm ratio, and high nucleus volume. Colours represent depth in the water column species are usually found, green = Pelagic; black = Benthopelagic; red = Demersal.

Habitat did not influence RBC shape (Pillai’s trace=0.6936; *P*=0.461). The results did not change if we use a multivariate PGLS (Table S4). We also found that evolution rate of RBC shape was very heterogeneous at the tip level (Fig. 5). The species with the highest rate was *Pygocentrus* (0.741) and the lowest was *Bujurquina* (0.087). There was a tendency to increase the rate in the common ancestor of *Pygocentrus, Metynnis*, and *Piaractus* in relation to the background rate, but the rate shift was not significant (rate difference = 0.392; *P* > 0.005). Interestingly, the species occupying the most extreme positions in the phylomorphospace, such as *Synbranchus* (0.514), *Astyanax* (0.555), and *Gymnotus* (0.17) were not the ones with the highest rates of evolution. Therefore, our results suggest that an increase in the rate of evolution does not necessarily produces morphological specialization and rate disparity is not the main pattern involved in the formation of RBC morphospace.

**Figure 5.**
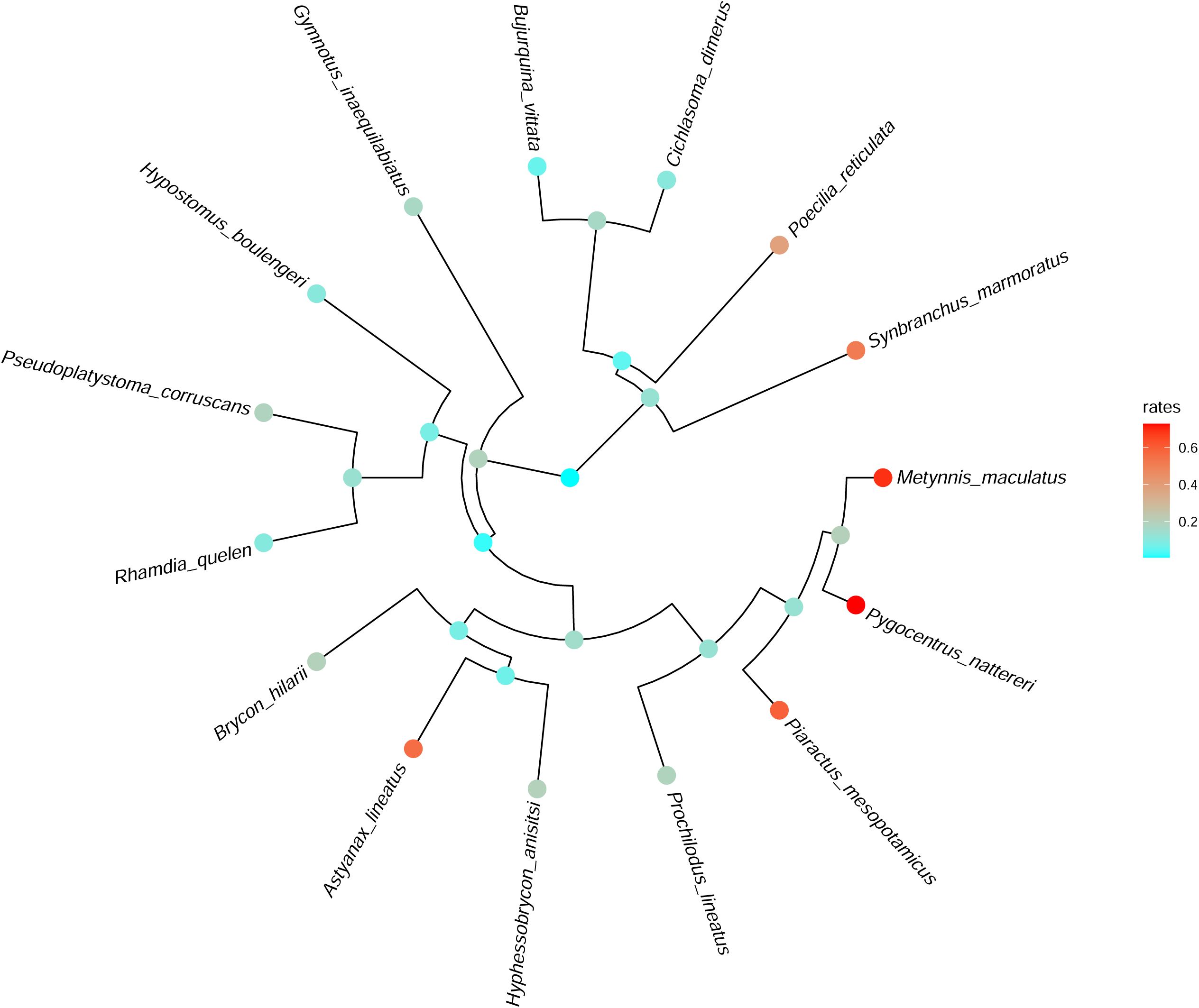
Phylogenetic tree showing the results of the tip-level evolution rate estimated using ridge regression for the stereological variables describing the Red Blood Cell shape. Analysis was conducted in RRphylo.

## Discussion

We found that the shape of Red Blood Cells (RBC) varies greatly among and within freshwater fish species. Contrarily to our initial hypothesis, we did not find a correlation between water column position (proxy for oxygen availability) and RBC shape. Accordingly, we did not find phylogenetic signal in RBC shape, suggesting closely related species do not have similar RBC shape. There does not seem to be a single mechanism driving the evolution of RBC shape, since the phylomorphospace shows a non-stationary pattern in the species similarity.

Small RBC allow species to occupy environments with low dissolved oxygen, since they usually have a higher number of cells, making them more efficient in oxygen uptake and transport (Silkin et al., 2019; Hawkey et al., 1991). Species with RBCs that have large surface area and volume have low metabolic rate, since the cell surface:volume ratio decreases the metabolic cost of gas exchange between membranes (Jones 1979, Lay and Baldwin, 1999). *Synbranchus* had higher perimeter (i.e., higher surface) and higher area. Thus, our results suggest that this species could have a lower metabolic rate than *Poecilia*, because the later has a smaller cell area and perimeter.

Most species occupied the center of the morphospace, independent of the phylogenetic relationship. This clustering of distantly related species suggests that convergence, apparently not caused by water column position, could be a mechanism influencing RBS shape evolution (Adams and Collyer 2019). Clustering in morphospace can also suggest a developmental constraint (Jablonski 2020) that impedes species to change their traits after speciation (Pie and Weitz 2005). Also, a few species were in extreme positions. Even though water column position did not explain RBC shape, the extreme position of *Synbranchus, Astyanax*, and *Poecilia* in the morphospace could be explained by differences in their life history strategies. For example, *Synbranchus* displays facultative pulmonary respiration, usually occurs in habitats with hypoxia, and burry itself during the dry season (Froese and Pauly, 2019). *Astyanax* also occurs in habitats with hypoxia (Froese and Pauly, 2019). The pattern we found points to clade disparity differences (Adams & Collyer 2019), with Acanthopterygii having higher dispersion in the morphospace than Ostariophysi (Fig. S6). These two orders have different ecological and biogeographic characteristics. Species from Ostariophysi are essentially from freshwater, while Acanthopterygii has marine origins, with secondary freshwater introgressions. One interesting pattern that emerges from the morphospace is that apparently there is no constrains in the morphospace, because almost all regions have one species. Siluriformes seem to have similar RBC shape, given their position in the morphospace, and similar evolution rate. This lineage occupies the upper position of the morphospace, along with *Piaractus* and *Hyphessobrycon*, and had high cell area and perimeter, and low nucleus:cytoplasm ratio, nucleus volume, and circularity. Exception for *Rhamdia* and *Hyphessobrycon*, all those species are demersal (benthic). Together these results suggest that the evolution of RBC shape was driven by fluctuating selection, although this result should be taken with care, since our sample size is not large.

We found a high heterogeneity in evolution rate of RBC shape. This high rate heterogeneity helps explain why we did not detect a significant phylogenetic signal, i.e., the change in shape seems to vary strongly at the tip level, instead of large clades. This is consistent with the model selection procedure that pointed Brownian Motion (BM) as best fit model to the data. BM can be caused by genetic drift, when selection randomly varies in direction through time, or when selection is weak compared to the time scale analysed (Felsenstein 1988). The large intraspecific variation in most variables (Table S1) may suggest that variables describing RBC shape are under weak selection pressure (Nikinmaa 2019). Interestingly, the position extreme positions of *Synbranchus* (0.514), *Astyanax* (0.555), and *Pygocentrus* (0.741) could be explained at least partially by an increase in evolution rate, which allowed them to diverge from their sister species, remaining Acanthopterygii, *Hyphessobrycon*, and *Metynnis* respectively. However, given our sample size we cannot rule out that RBC shape of these species are evolving under directional selection. Conversely, if lineages followed a random walk (akin of BM) one would expect species to diverge and occupy positions in the morphospace irrespective of their habitat or phylogenetic position (Pie and Weitz 2005).

Water column position did not explain RBC shape in this freshwater fish species radiation. This result suggest that RBC shape may not be entirely determined by partial pressure of dissolved oxygen in water. It also suggests that water column position does not represent a main constraint in the evolution of these traits. A recent study (Minias 2020) also found that life history traits did not explain hemoglobin and hematocrit evolution in birds. Differently from mammals, fish RBC are nucleated cells and have many cytoplasmic organelles, such as mitochondria, Golgi complex, centrioles, ribosomes, microtubules, rough and smooth endoplasmic reticulum (Sekhon and Beams 1969), as well as a range of cytoplasmatic enzymes involved in the anaerobic glycolysis (Sephton et al. 1991). These organelles and enzymes allow the functions of RBC to be finely regulated and somewhat independent from other components of the hematopoietic system, enabling fish to occupy a wide range of environments (Nikinmaa et al. 2019). Therefore, species may respond rapidly to changes in oxygen availability (Witeska, 2013), which indicates that it is not constrained by inheritance (Arnold 1992) or phylogenetic inertia, at least to a certain degree.

In conclusion, our study provides a novel macroevolutionary perspective on fish RBC shape by combining commonly used design-based stereological techniques with modern phylogenetic comparative methods. We found that the RBC of the two main orders of Neotropical freshwater fish vary greatly in shape, but that their water column alone does not explain it. Future studies should use quantitative genetic approaches to explore the underlying genetic basis for RBC morphology, possibly using geometric morphometrics and experimental approaches motivated by ecological data.

## Acknowledgements

This study was supported by Universidade Federal de Mato Grosso do Sul – UFMS/MEC – Brasil. This study was funded in part by the Coordenação de Aperfeiçoamento de Pessoal de Nível Superior – Brasil (CAPES) – Finance Code 001. F. R. Carvalho and K. Tondato provided insightful comments as members of BOM bachelor’s honors thesis examining committee.

## Author contributions

BOM collected cell morphometric data assisted by MSS, LFB. CEF and DBP conceived the study. DBP analysed and curated the data. DBP, LFB, and BOM prepared the first draft of the manuscript to which all authors contributed afterwards.

## Data accessibility statement

All the data and R code used to run the analysis will be deposited in FigShare upon acceptance.

