## Supplementary material for "The evolution of red blood cell shape in a continental radiation of fishes"

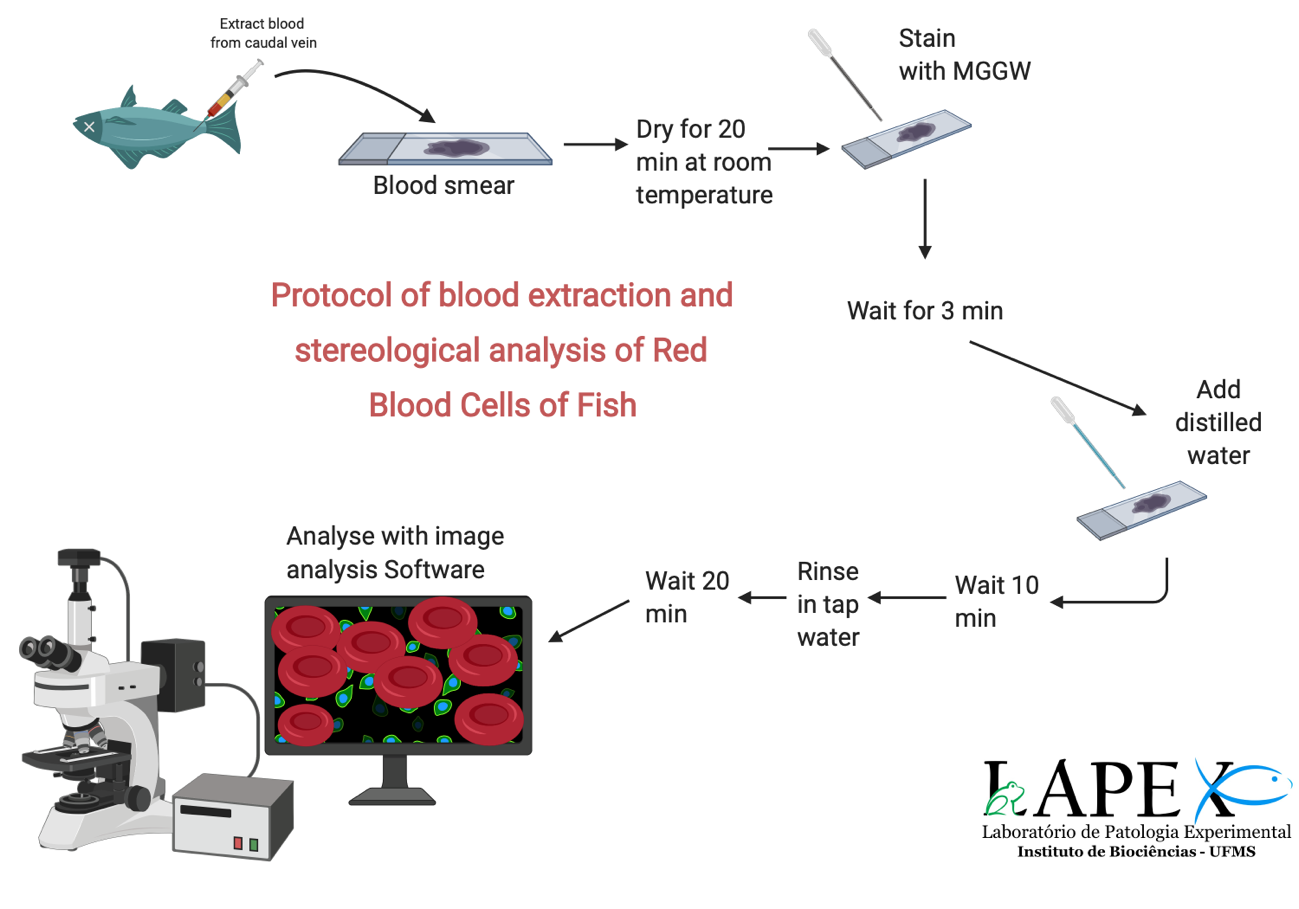

Figure S1. Summary of the protocol for blood extraction of fish and stereological analysis of Red Blood Cells. The full protocol is available at https://protocols.io/view/staining-of-fish-red-blood-cells-bd9yi97w.html

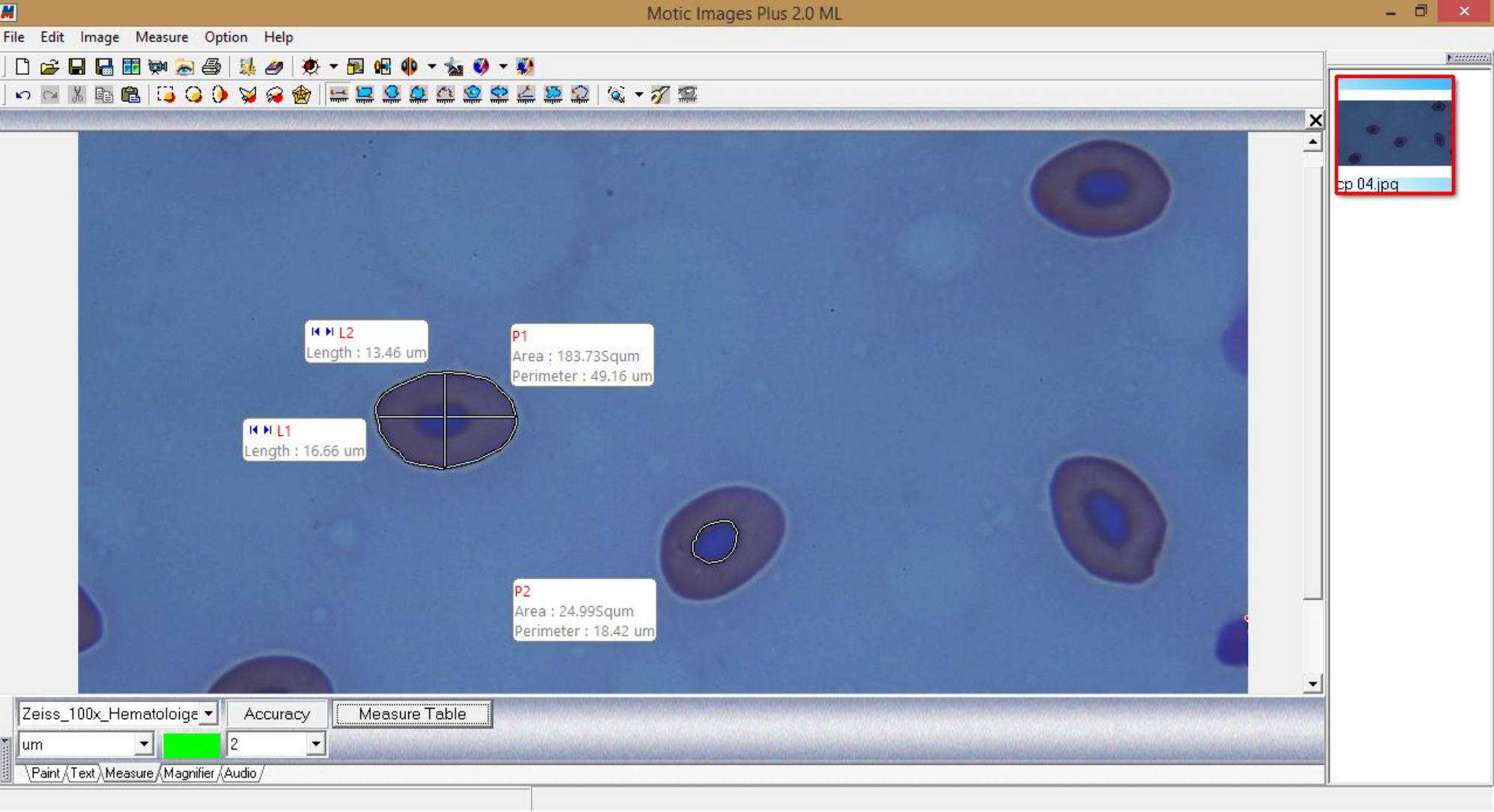

Figure S2. Screen capture of the software we used to measure stereological variables, showing how linear (first-order) measurements were taken. P1 means circular selection, which gives the perimeter (µm) and area (µm^2^) of the cell, the same for the nucleus (P2). L1 is the largest diameter, while L2 is the smallest diameter (µm).

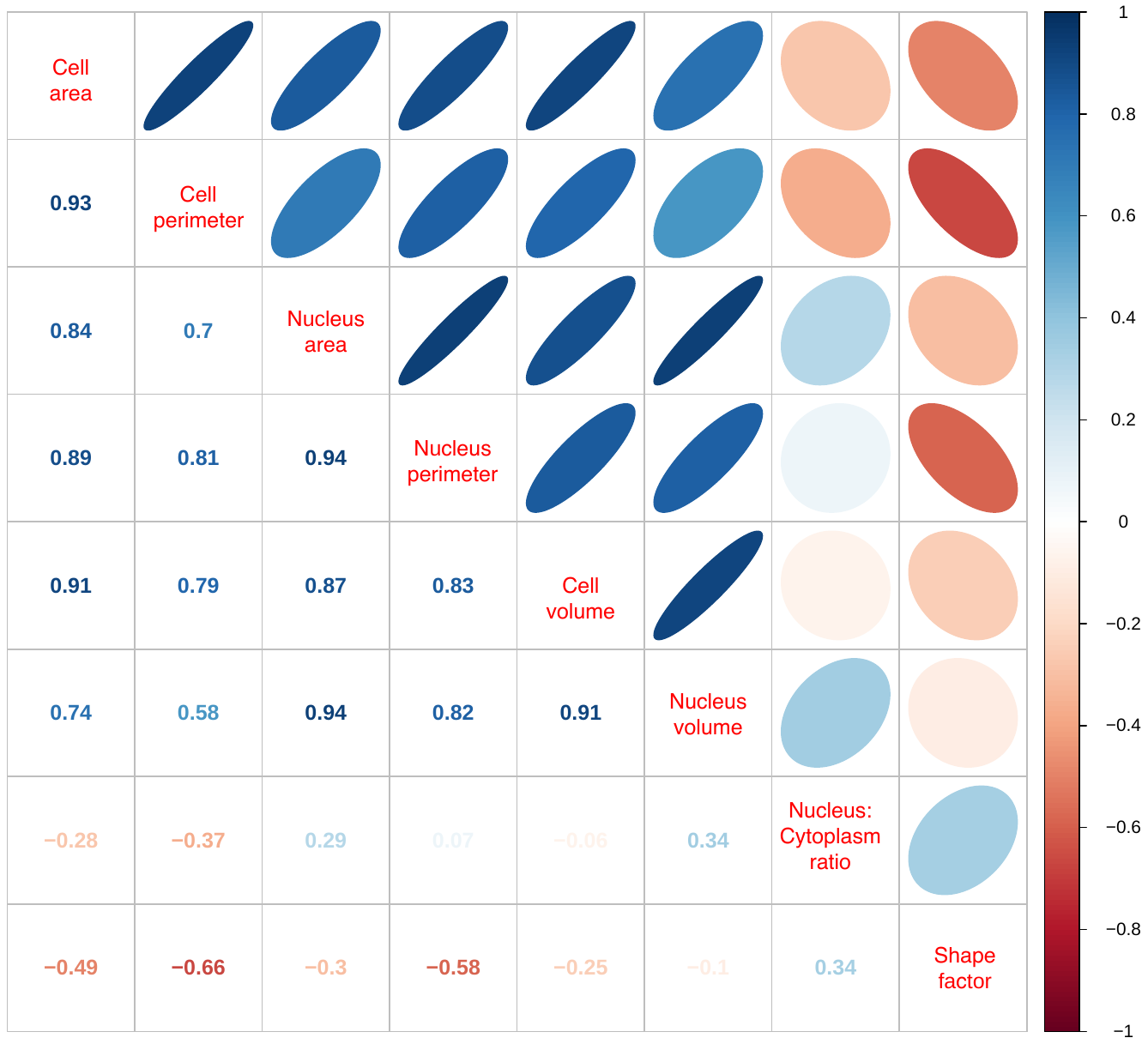

Figure S3. Correlogram showing the pair-wise correlation structure between first-order and second-order stereological variables calculated using the mean values for all species.

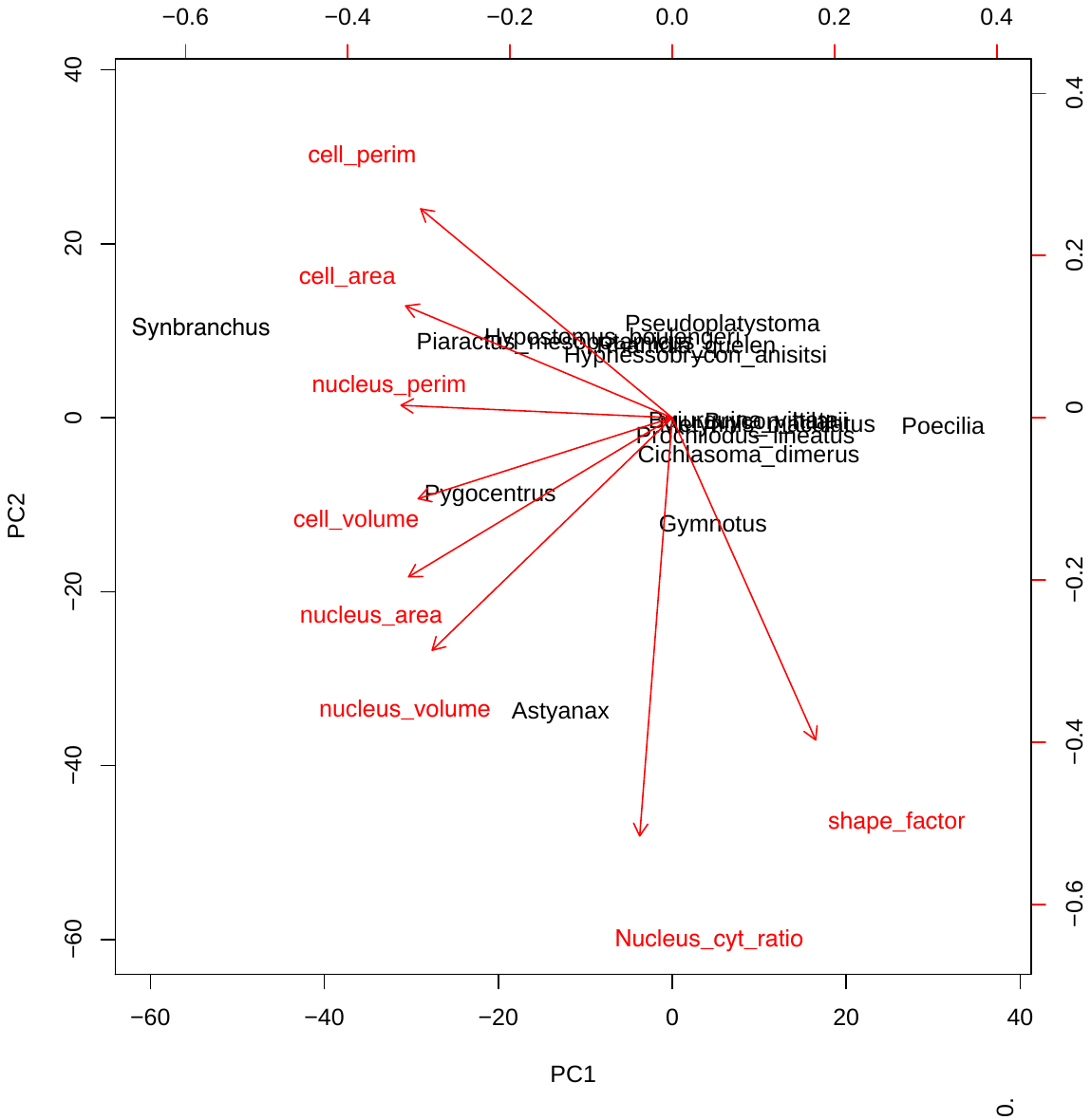

Figure S4. Ordination diagram showing the result of phylogenetic Principal Components Analysis. Stereological variables are in red and species are in black.

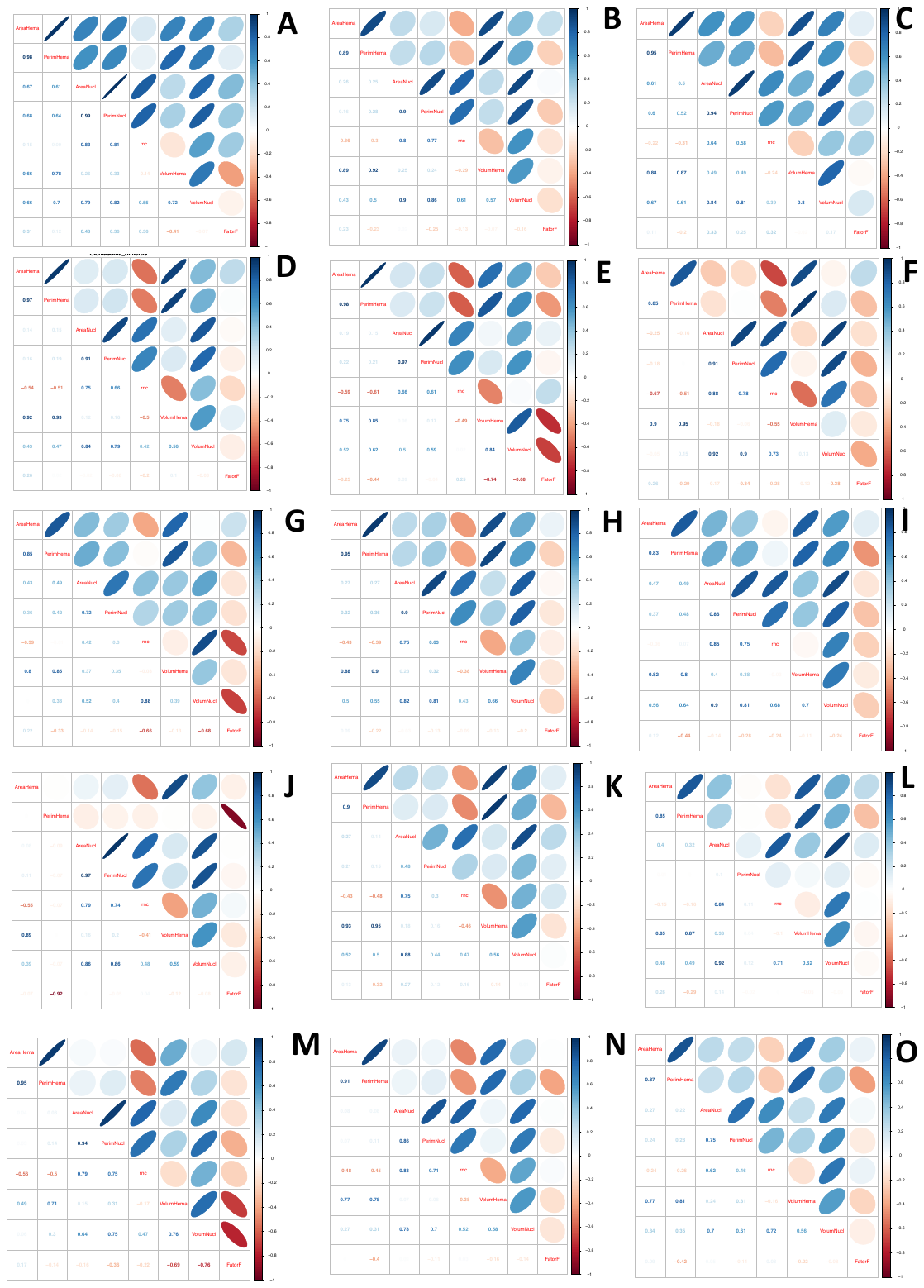

Figure S5. Phenotypic matrices (**P** matrix) of each species showing the change in covariance structure among variables. A) *Astyanax*, B) *Brycon*, C) *Bujurquina*, D) *Cichlasoma*, E) *Gymnotus*, F) *Hyphessobrycon*, G) *Hypostomus*, H) *Metynnis*, I) *Piaractus*, J) *Poecilia*, K) *Prochilodus*, L) *Pseudoplatystoma*, M) *Pygocentrus*, N) *Rhamdia*, O) *Synbranchus*.

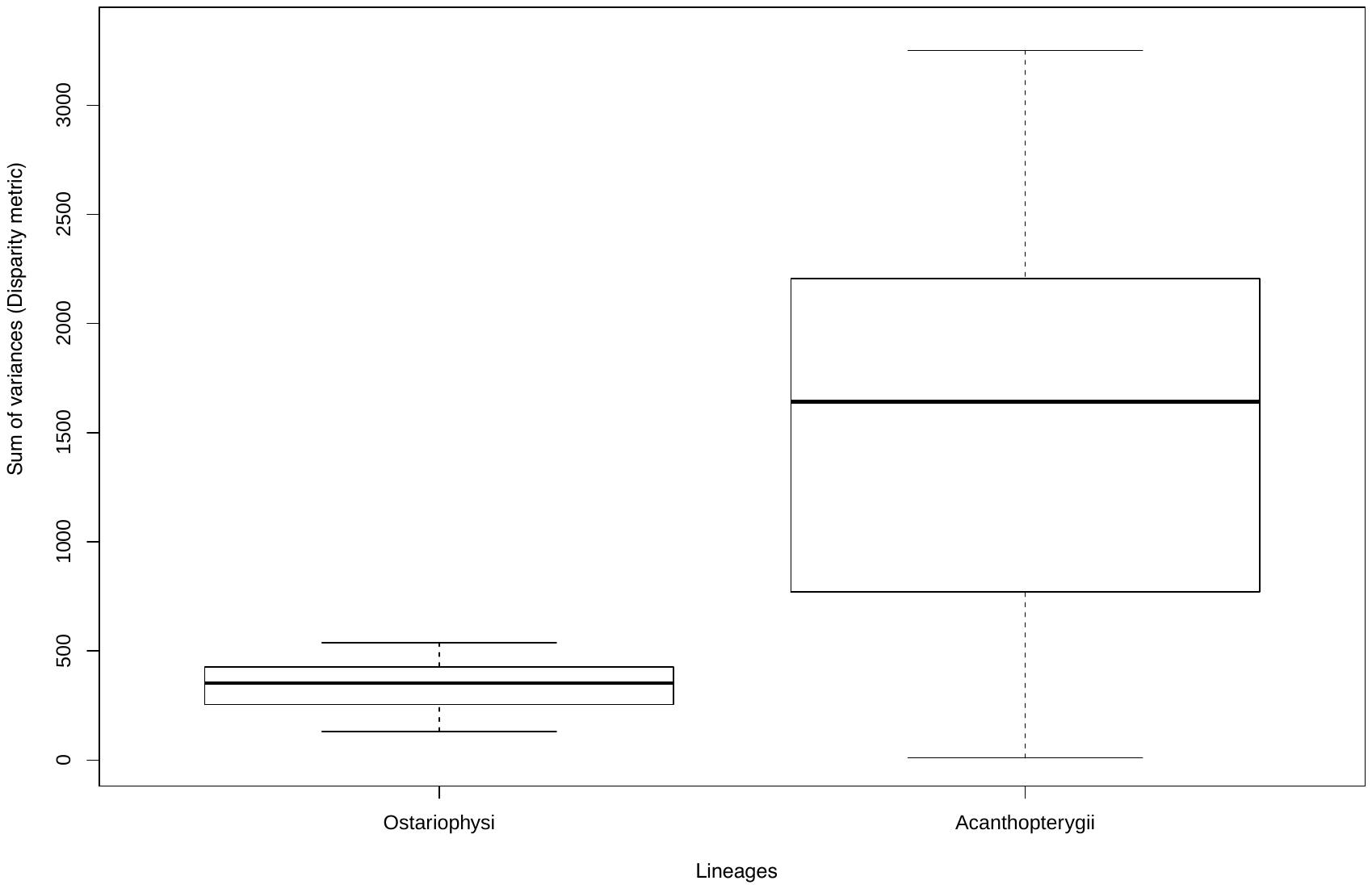

Figure S6. Boxplot showing the differences in morphological disparity between the two superorders Ostariophysi and Acanthopterygii calculated using the sum of variances. Differences in disparity are significant after a Wilcoxon rank test (W = 7729; *P* < 0.0001). Result does not change if we use alternative disparity metrics. Analysis was run using the dispRity R package (Guillerme 2018 Methods in Ecology and Evolution).

Table S1. Mean and standard deviation of stereological variables of Red Blood Cells of 15 freshwater fish species. Measurements were taken in 250 cells per species (50 cells per specimen * 5 specimens). Cell and nucleus area and perimeter are first-order stereological variables, while the others are second-order variables.

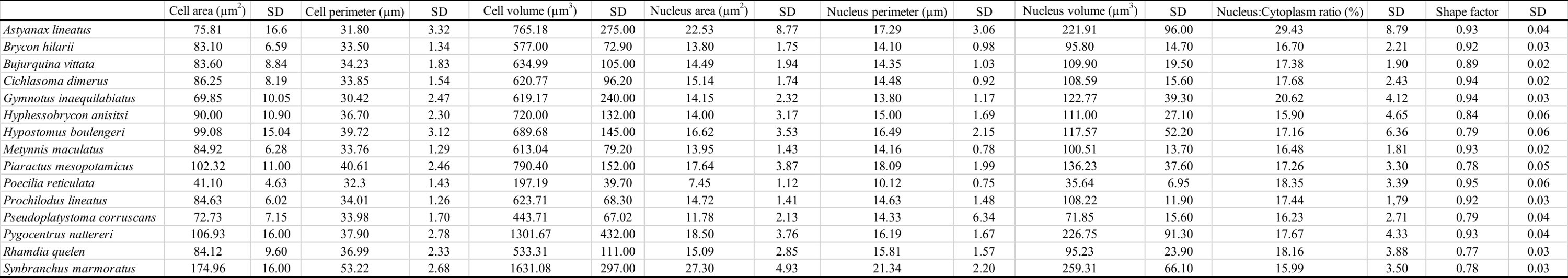

Table S2. Results of the phylogenetic Principal Components Analysis showing the loadings of stereological variables describing the shape of Red Blood Cells. Bold face values indicate high correlation (>0.80).

|  | pPC1 | pPC2 | pPC3 |
| --- | --- | --- | --- |
| Cell area | **-0.9335399** | 0.23344805 | 0.202862399 |
| Cell perimeter | **-0.8808330** | 0.43603175 | 0.006961059 |
| Nucleus area | **-0.9230393** | -0.33214086 | -0.122118950 |
| Nucleus perimeter | **-0.9498726** | 0.02596405 | -0.278039656 |
| Nucleus:cytoplasm ratio | -0.1138561 | **-0.87313302** | -0.468062859 |
| Cell volume | **-0.8896341** | -0.16902536 | 0.388276950 |
| Nucleus volume | **-0.8405402** | -0.48540674 | 0.172702041 |
| Shape factor (circularity) | 0.5020555 | -0.67263151 | 0.509872763 |
| Relative Eigenvalue (%) | 64.6 | 22.9 | 9.9 |

Table S3. Results of the evolutionary model selection conducted in mvMORPH for the Red Blood Cell shape (described by the eight phylogenetic Principal Components). EIC means Extended Information Criterion. Bold face indicates the unequivocal best fit model.

| Model | EIC | ΔEIC | Standard Error | Log likelihood |
| --- | --- | --- | --- | --- |
| Pagel’s λ | 780.8582 | 14.3587 | 108.9735 | -256.4494 |
| Brownian Motion | **766.4995** | **0** | 82.46245 | -275.952 |
| Ornstein-Uhlenbeck | 803.6616 | 37.1621 | 105.8629 | -255.629 |

Table S4. Results of the Phylogenetic Generalized Least Squares (PGLS) using Type I Sum of Squares and residual randomization conducted in geomorph to test the effect of habitat on Red Blood Cell shape (described by the eight pPCs). The model assumes implicitly that traits have evolved under a Brownian Motion model of evolution.

|  | DF | SS | MS | R^2^ | F | Z | *P* |
| --- | --- | --- | --- | --- | --- | --- | --- |
| Habitat | 2 | 15.570 | 7.7848 | 0.14272 | 0.9989 | 0.28205 | 0.402 |
| Residuals | 12 | 93.522 | 7.7935 | 0.85728 |  |  |  |
| Total | 14 | 109.092 |  |  |  |  |  |
